# Deep learning-based cross-classifications reveal conserved spatial behaviors within tumor histological images

**DOI:** 10.1101/715656

**Authors:** Javad Noorbakhsh, Saman Farahmand, Ali Foroughi pour, Sandeep Namburi, Dennis Caruana, David Rimm, Mohammad Soltanieh-ha, Kourosh Zarringhalam, Jeffrey H. Chuang

## Abstract

Histopathological images are a rich but incompletely explored data type for studying cancer. Manual inspection is time consuming, making it challenging to use for image data mining. Here we show that convolutional neural networks (CNNs) can be systematically applied across cancer types, enabling comparisons to reveal shared spatial behaviors. We develop CNN architectures to analyze 27,815 hematoxylin and eosin slides from The Cancer Genome Atlas for tumor/normal, cancer subtype, and mutation classification. Our CNNs are able to classify tumor/normal status of whole slide images (WSIs) in 19 cancer types with consistently high AUCs (0.995±0.008), as well as subtypes with lower but significant accuracy (AUC 0.87±0.1). Remarkably, tumor/normal CNNs trained on one tissue are effective in others (AUC 0.88±0.11), with classifier relationships also recapitulating known adenocarcinoma, carcinoma, and developmental biology. Moreover, classifier comparisons reveal intra-slide spatial similarities, with average tile-level correlation of 0.45±0.16 between classifier pairs. Breast cancers, bladder cancers, and uterine cancers have spatial patterns that are particularly easy to detect, suggesting these cancers can be canonical types for image analysis. Patterns for TP53 mutations can also be detected, with WSI self- and cross-tissue AUCs ranging from 0.65-0.80. Finally, we comparatively evaluate CNNs on 170 breast and colon cancer images with pathologist-annotated nuclei, finding that both cellular and intercellular regions contribute to CNN accuracy. These results demonstrate the power of CNNs not only for histopathological classification, but also for cross-comparisons to reveal conserved spatial biology.

## Introduction

Histopathological images are a crucial data type for diagnosis of cancer malignancy and selecting treatment (He et al. 2012), indicative of their value for understanding cancer biology. However, manual analysis of whole slide images (WSIs) is labor-intensive (Gurcan et al. 2009) and can vary by observer (Allison et al. 2014; Stang et al. 2006; Grilley-Olson et al. 2013), making it difficult to scale such approaches for discovery-oriented analysis of large image collections. Image datasets for hematoxylin and eosin (H&E), immunohistochemistry (IHC), and spatial -omic imaging technologies are rapidly growing (Litjens et al. 2017). Improved computational approaches for analyzing cancer images would therefore be valuable, not only for traditional tasks such as histopathological classification and cell segmentation (Cooper et al. 2018), but also for novel questions such as the de novo discovery of spatial patterns that distinguish cancer types. The search for recurrent spatial patterns is analogous to the search for common driver mutations or expression signatures based on cancer sequencing (Bailey et al. 2018), yet this paradigm has been little explored for cancer image data.

In the last few years, there have been major advances in supervised and unsupervised learning in computational image analysis and classification (Russakovsky et al. 2015; Litjens et al. 2017), providing opportunities for application to tumor histopathology. Manual analysis involves assessments of features such as cellular morphology, nuclear structure, or tissue architecture, and such pre-specified image features have been inputted into support vector machines or random forests for tumor subtype classification and survival outcome analysis, e.g. (Luo et al. 2017; Yu et al. 2016; Mousavi et al. 2015). However, pre-specified features may not generalize well across tumor types, so recent studies have focused on fully-automated approaches using convolutional neural networks (CNNs), bypassing the feature specification step. For example, Schaumberg et. al., trained ResNet-50 CNNs to predict SPOP mutations using WSIs from 177 prostate cancer patients (Schaumberg, Rubin, and Fuchs 2018), achieving AUC = 0.74 in cross validation and AUC = 0.64 on an independent cohort. Yu et al., utilized CNN architectures including AlexNet, GoogLeNet, VGGNet-16 (Simonyan and Zisserman 2014), and the ResNet-50 to identify transcriptomic subtypes of lung adenocarcinoma (LUAD) and squamous cell carcinoma (LUSC) (Yu et al. 2019). They were able to classify LUAD vs. LUSC (AUC of 0.88-0.93), as well as each vs. adjacent benign tissues with higher accuracy. Moreover, they were able to predict the TCGA transcriptomic classical, basal, secretory, and primitive subtypes of LUAD (Wilkerson et al. 2010, 2012) with AUCs 0.77-0.89, and similar subtype classifications have been reported in breast (Couture et al. 2019). Recently, Coudray et al. (Coudray et al. 2018) proposed a CNN based on Inception v3 architecture to classify WSIs in LUAD and LUSC, achieving an AUC of in tumor/normal classification. Further, their models were able to predict mutations in 10 genes in LUAD with AUCs 0.64-0.86, and subsequently mutations in BRAF (AUC ∼ 0.75) or NRAS (AUC ∼ 0.77) melanomas (Kim et al. 2019). Other groups have used CNNs to distinguish tumors with high or low mutation burden (Xu et al. 2019). These advances highlight the potential of CNNs in computer assisted analysis of WSIs.

Many critical questions remain. For example, prior studies have focused on individual cancer types, but there has been little investigation of how neural networks trained on one cancer type perform on other cancer types, which could provide important biological insights. As an analogy, comparisons of sequences from different cancers have revealed common driver mutations (Martincorena et al. 2017; Hoadley et al. 2018), e.g. both breast and gastric cancers have frequent HER2 amplifications, and both are susceptible to treatment by trastuzumab (Bang et al. 2010; Piccart-Gebhart et al. 2005). Such analysis is in a rudimentary state for image data, as it remains unclear how commonly spatial behaviors are shared between cancer types. A second important question is the impact of transfer learning on cancer image analysis. Transfer learning is used to pre-train neural networks using existing image compilations (Zhou et al, 2011). However, standard compilations are not histological, and it is unclear how this affects cancer studies. A third key topic is to clarify the features that impact prediction accuracy. For example, recurrent neural network approaches (Campanella et al., 2019) have been shown to distinguish prostate, skin and breast cancers at the slide level, but the relevant spatial features are not well understood. Determination of predictive features is affected not only by the underlying biology, but also by availability of spatial annotations and appropriate computational techniques.

To investigate these questions, here we analyze 27,815 frozen or FFPE whole-slide H&E images from 23 cohorts from The Cancer Genome Atlas (TCGA), a resource with centralized rules for image collection, sequencing and sample processing. We have developed image processing and convolutional neural network software that can be broadly applied across tumor types to enable cross-tissue analyses. Using these techniques, first, we show that this CNN architecture can distinguish tumor/normal and cancer subtypes in a wide range of tissue types. Second, we systematically compare the ability of neural networks trained on one cancer type to classify images from another cancer type. We show that cross-classification relationships recapitulate known tissue biology. Remarkably, these comparisons also reveal that breast, bladder, and uterine cancers can be considered canonical cancer image types. Third, we investigate driver effects by determining how cancers with the TP53 mutation can be cross-classified across tissues, including a comparison of transfer learning vs. full CNN training. Fourth, we test how cellular vs. intercellular regions impact CNN tumor/normal predictions, making use of cell-resolution annotations from 170 colorectal and breast cancer images. Our studies demonstrate that cross-comparison of CNN classifiers is a powerful approach for discovering shared biology within cancer images.

## Results

### Pan-cancer convolutional neural networks for tumor/normal classification

We developed a CNN architecture to classify slides from TCGA by tumor/normal status, using a neural network that feeds the last fully connected layer of an Inception v3-based CNN pre-trained on ImageNet into a fully connected layer with 1024 neurons. This architecture is depicted in Figure 1a, and a related architecture for mutation classification (described in sections below) is shown for comparison in Figure 1b. The two final fully connected layers of the tumor/normal CNN were trained on tiles of size 512×512 from WSIs. We trained this model separately on flash frozen slides from 19 TCGA cohorts having numbers of slides ranging from 205-1,949 (Figure 2a). 70% of the slides were randomly assigned to the training set and the rest were assigned to the test set. To address the data imbalance problem (Charte et al. 2015), the majority class was undersampled to match the minority class.

**Figure 1.**
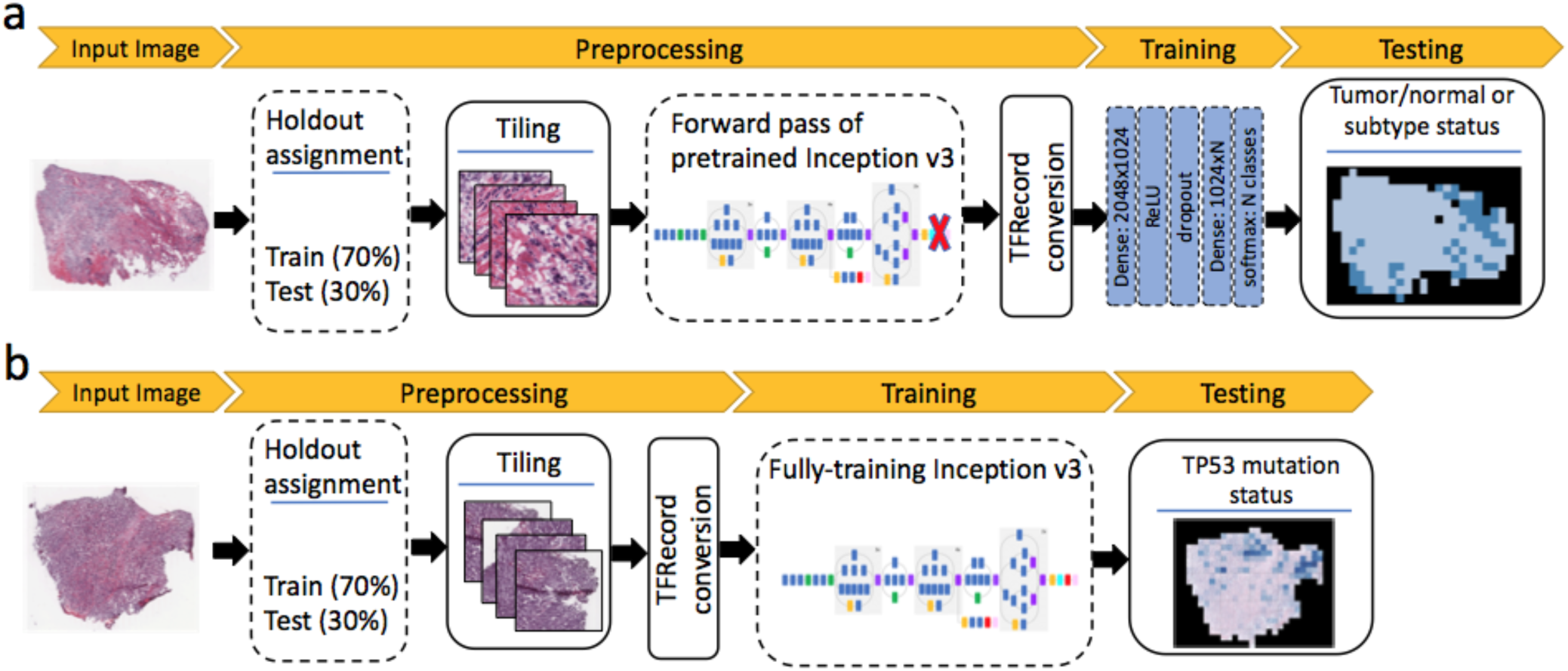
Classification pipelines. a) Transfer learning pipeline for tumor/normal and subtype classification b) Full training pipeline for mutation classification

**Figure 2.**
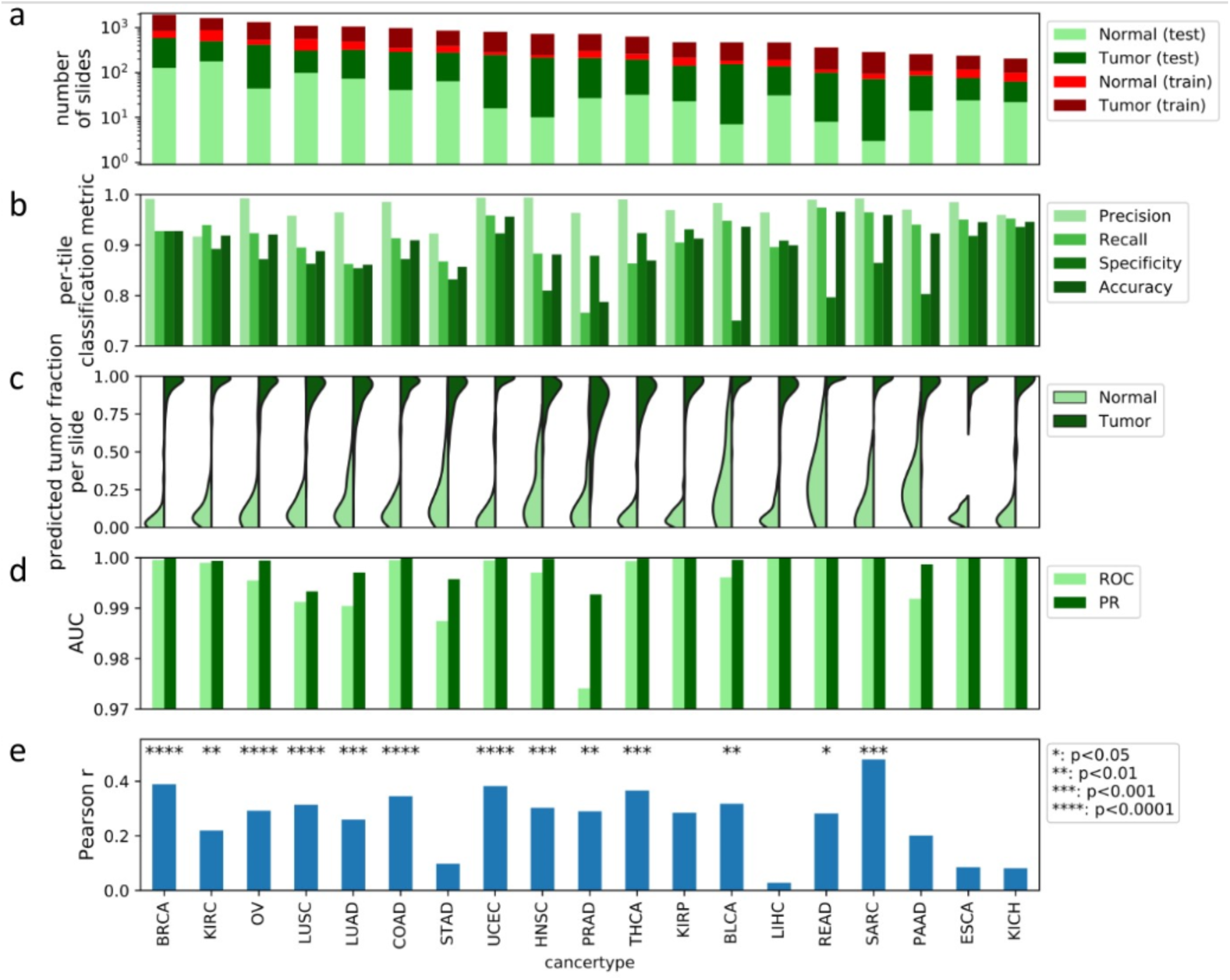
Tumor/normal classification using CNNs. (a) Numbers of tumor and normal slides in test and training sets. (b) Per-tile classification metrics. (c) Fraction of tiles within each slide predicted as tumor. (d) per slide AUC values for tumor/normal classification for ROC and precision-recall curve. (PR) (e) Pearson correlation coefficients between predicted and pathologist evaluation of tumor purity. P-values are based on permutation test of the dependent variable after Bonferroni correction across all cohorts.

Figure 2b shows the classification results. We used a naive training approach such that, after removal of background regions, all tiles in a normal image are assumed normal and all tiles in a tumor image are assumed tumor. The CNN accurately classifies test tiles for most tumor types (accuracy: 0.91±0.05, precision: 0.97±0.02, recall: 0.90±0.06, specificity: 0.86±0.07. Mean and standard deviation calculated across cohorts). We next examined the fraction of tiles classified as tumor or normal within each slide. The fractions of tiles matching the slide annotation are 0.88±0.14 and 0.90±0.13 for normal and tumor samples, respectively (Figure 2c) (mean and standard deviation calculated from all cohorts pooled together). These fractions are high in almost all slides, and the tumor-predicted fraction (TPF) is significantly different between tumors and normals (p < 0.0001 per-cohort comparison of tumor vs. normal, Welch’s t-test). We also performed the classification on a per-slide basis. To do this, we used the TPF in each slide as a metric to classify it as tumor or normal. This approach yielded extremely accurate classification results for all cohorts (Figure 2d, mean AUC ROC = 0.995, mean PR AUC = 0.998). Confidence intervals (CI) of per-slide predictions are given in Figure S1 (also see Methods). The CI lower bound on all classification models was above 90%, with cancer types having fewer slides or imbalanced test data tending to have larger CIs. These results indicate that our network can successfully classify WSIs as tumor/normal across many different cancer types.

We next investigated whether TPF is a meaningful measure of tumor purity. We found significant positive correlations between TPF and TCGA pathologist-reported purity in the majority of cancer types (Figure 2e), with larger cohorts tending to have more significant p-values (e.g. BRCA: p=5e-17). The distributions of TPF were systematically higher than the pathologist annotations (Figure S2), though this difference can be reconciled by the fact that TPF is based on neoplastic area while the pathologist annotation is based on cell counts. Tumor cells are larger than stromal cells and reduce the nuclear density. While TPF and purity are clearly related, the moderate magnitudes of correlations indicate that intra-slide improvements can be made. A notable limitation is the training assumption that tiles in a slide are either all tumor or all normal, as intra-slide pathologist annotations are not provided by TCGA.

### Neural network classification of cancer subtypes

We also applied our algorithm to classify tumor slides based on their cancer subtypes (Figure 1a). This analysis was performed on 10 tissues for which pathologist subtype annotation was available on TCGA: sarcoma (SARC), brain (LGG), breast (BRCA), cervix (CESC), esophagus (ESCA), kidney (KIRC/KIRP/KICH), lung (LUAD/LUSC), stomach (STAD), uterine (UCS/UCEC), and testis (TGCT). Cancer subtypes with at least 15 samples were considered, based on TCGA metadata (see Methods). Because comparable numbers of FFPE and flash frozen samples are present in TCGA cohorts (FFPE to frozen slide ratio: 1.0±0.5), both were included (Figure 3a). The following subtypes were considered (Figure 3b): brain (oligoastrocytoma, oligodendroglioma, astrocytoma), breast (mucinous, mixed, lobular, ductal), cervix (adenoma, squamous cell carcinoma), esophagus (adenocarcinoma, squamous cell carcinoma), kidney (chromophobe, clear cell, papillary), lung (adenocarcinoma, squamous cell carcinoma), sarcoma (MFS: myxofibrosarcoma, UPS: undifferentiated pleomorphic sarcoma, DDLS: dedifferentiated liposarcoma, LMS: leiomyosarcomas), stomach (diffuse, intestinal), testis (non-seminoma, seminoma), thyroid (tall, follicular, classical), uterine (carcinoma, carcinosarcoma). We used the same CNN model as for tumor/normal classification; however, for cohorts with more than two subtypes, a multi-class classification was used.

**Figure 3.**
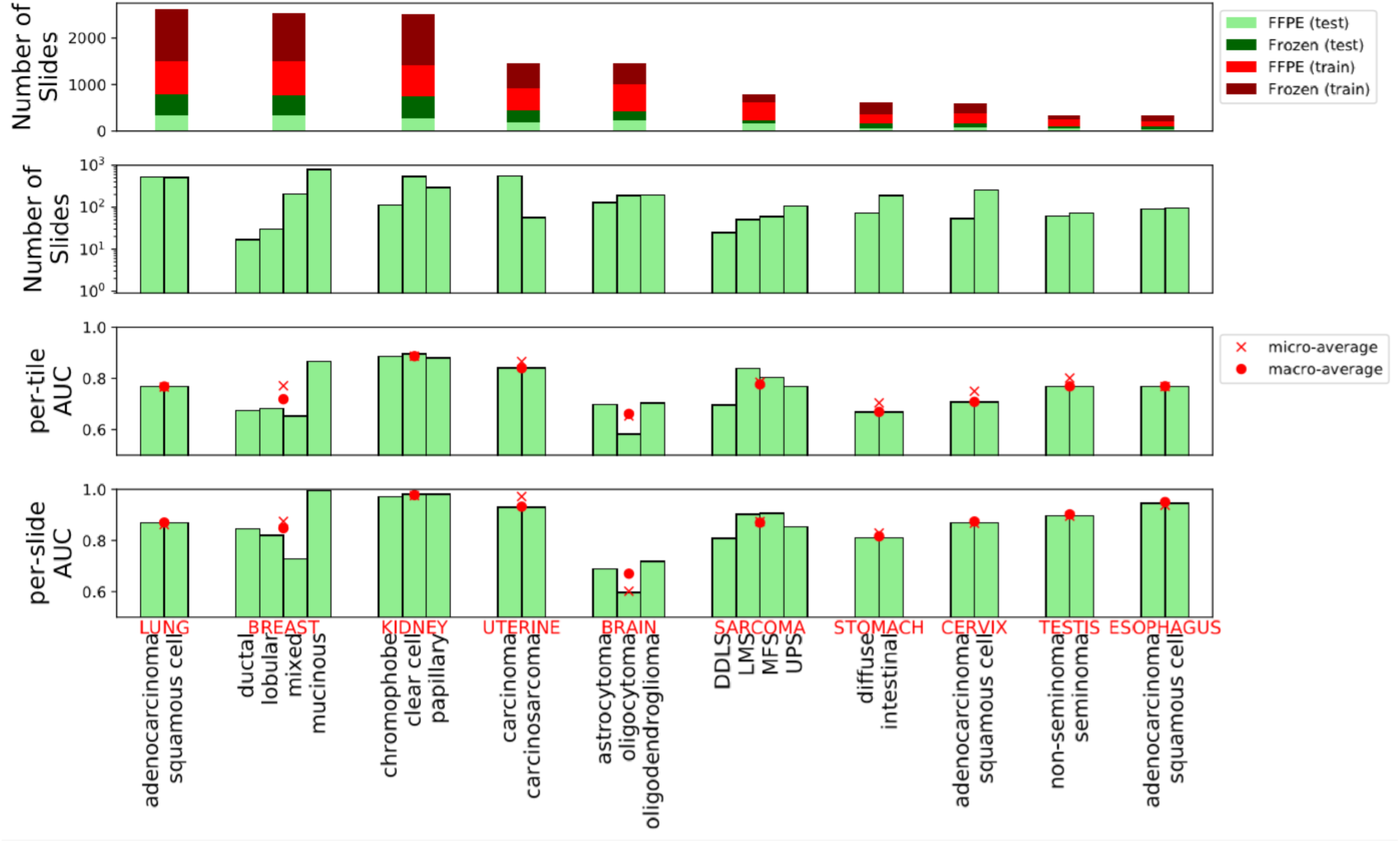
Subtype classification using CNNs. (a) Number of samples used for training. (b) Number of samples for each subtype. (c) AUC ROC for subtype classifications at the tile level (d) and at the slide level.

Figure 3c and 3d show the per-tile and per-slide classification results (AUC ROCs alongside their micro- and macro-averages). At the slide level, the classifiers can identify the subtypes with good accuracy in most tissues, though generally not yet at clinical precision (AUC micro-average: 0.87±0.1; macro-average: 0.87±0.09). The tissue with the highest AUC micro/macro-average was kidney (AUC 0.98), while the lowest was brain with micro-average 0.60 and macro-average 0.67. All CIs were above the 0.50 null AUC expectation, and all of the AUCs were statistically significant (5% FDR, Benjamini-Hochberg correction (Benjamini and Hochberg, 1995)). For full CIs and p-values, see Table S1. The individual subtype with best AUC is the mucinous subtype for breast cancer (adjusted p-value <1e-300). The weakest p-value (adjusted p=0.012) belongs to the oligoastrocytoma subtype of the brain. Slide predictions are superior to those at the tile level, though with similar trends across tissues. This indicates that tile averaging provides substantial improvement of signal to noise, consistent with observations for the tumor/normal analysis.

### Cross-classifications between tumor types demonstrate conserved spatial behaviors

We next used cross-classification to test the hypothesis that different tumor types share CNN-detectable morphological features distinct from those in normal tissues. For each cancer cohort, we re-trained the binary CNN classifier for tumor/normal status using all WSIs in the cohort. We then tested the ability of each classifier to predict tumor/normal status in the samples from each other cohort. Figure 4 shows a heat map of per-slide AUC for all cross-classifications, hierarchically clustered on the rows and columns of the matrix. A non-clustered version is presented in Figure S3 with CIs. Surprisingly, neural networks trained on any single tissue were successful in classifying cancer vs. normal in most other tissues (average pairwise AUCs of off-diagonal elements: 0.88±0.11 across all 342 cross-classifications). This prevalence of strong cross-classification supports the existence of morphological features shared across cancer cohorts but not normal tissues. In particular, classifiers trained on most cohorts successfully predicted tumor/normal status in BLCA (AUC=0.98±0.02), UCEC (AUC=0.97±0.03), and BRCA (AUC=0.97±0.04), suggesting that these cancers most clearly display features universal across types. At a 5% FDR, 330 cross-classification AUCs are significant (See Figure S3 for statistical details). The AUC mean and CI lower bound are each above 80% for 300 and 164 of these cross-classifications, respectively. A few cancer types, e.g. LIHC and PAAD, showed poor cross-classification to other tumor types, suggesting morphology distinct from other cancers.

**Figure 4.**
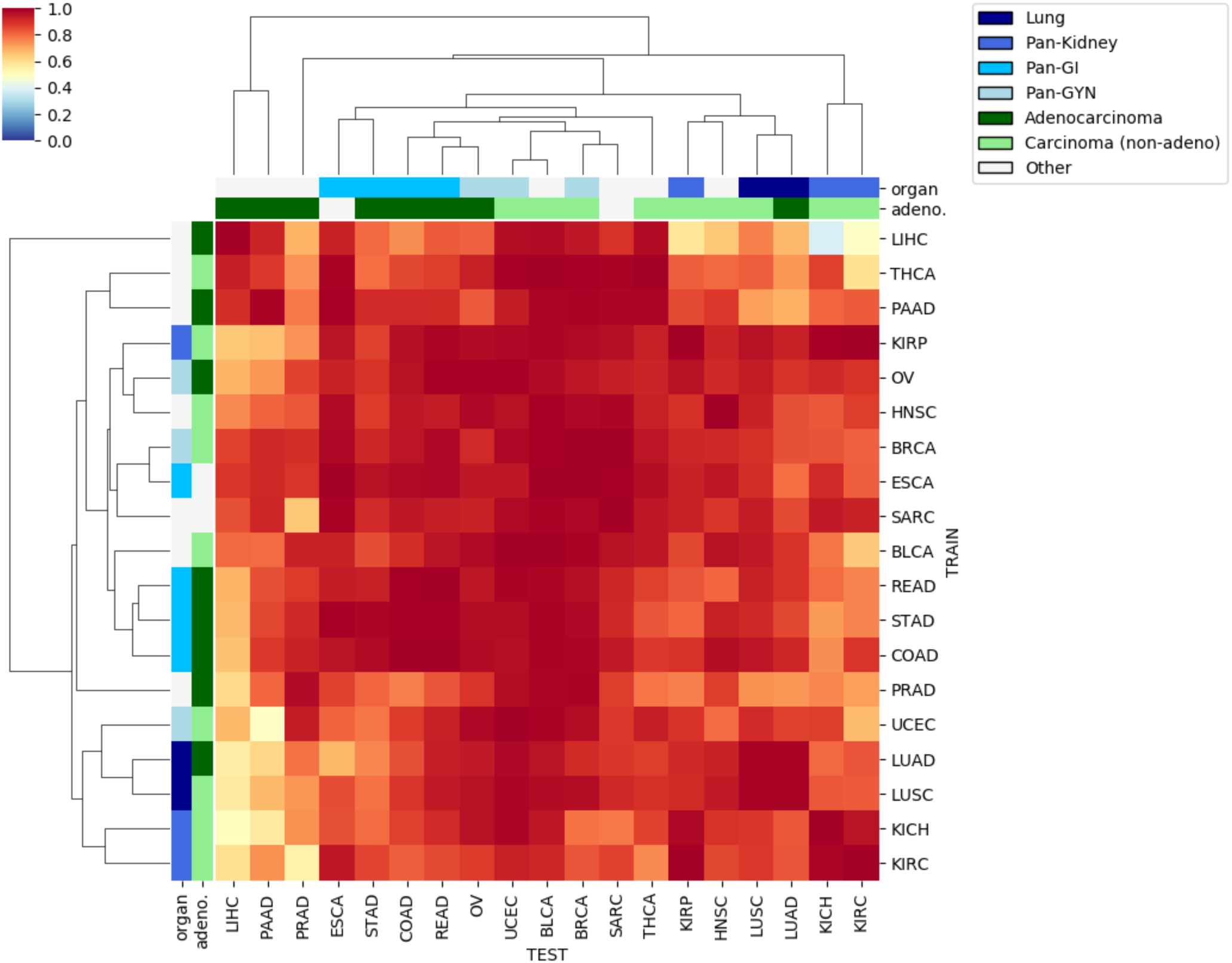
Per-slide AUC values for cross classification of tumor/normal status. The hierarchically clustered heatmap shows pairwise AUC values of CNNs trained on the tumor/normal status of one cohort (train axis) tested on the tumor/normal status of another cohort (test axis). Adeno-ness (adenocarcinoma vs. non-adeno carcinoma) and organ of origin (lung, kidney, gastrointestinal, gynecological) for each cohort are marked with colors on the margins. Cohorts with ambiguous or mixed phenotype are marked as ‘Other’.

To improve spatial understanding of these relationships, we tested how well tile-level predictions are conserved between different classifiers (Figure 5), while also analyzing the effect of varying the test set. For each pair of classifiers, we specified a test set then computed the correlation coefficient of the predicted tumor/normal state (logit of the tumor probability) across all tiles in the test set. We repeated this calculation for each test set, which we indexed by tissue type (breast, bladder, etc.). Each test set included both tumor and normal slides for the tissue type. Figures 5a and 5b show for each pair of classifiers the average and maximum correlation coefficients, respectively, over test sets. Many correlations are positive, with an average and standard deviation over all pairs of classifiers of 45±16% (Fig. 5a, diagonal elements excluded), indicating cross-classifiers agree at the tile level. These tile-level results supported the slide-level results. Classifiers with low cross-classification slide-level AUCs, such as LIHC, had the smallest tile level correlations. Tile predictions also showed similarities between classifiers derived from the same tissue (e.g. LUAD-LUSC, KICH-KIRP-KIRC). Similarities between classifiers became even more apparent when we focused on the test tissue with the strongest correlation for each pair of classifiers (Figure 5b). These positive correlations are not simply due to distinguishing tiles in tumor slides from tiles in normal slides. Figures 5c and 5d are analogous to Figures 5a and 5b, but computed only over tumor slides. The results are nearly unchanged, indicating that they reflect biology within tumor images.

**Figure 5.**
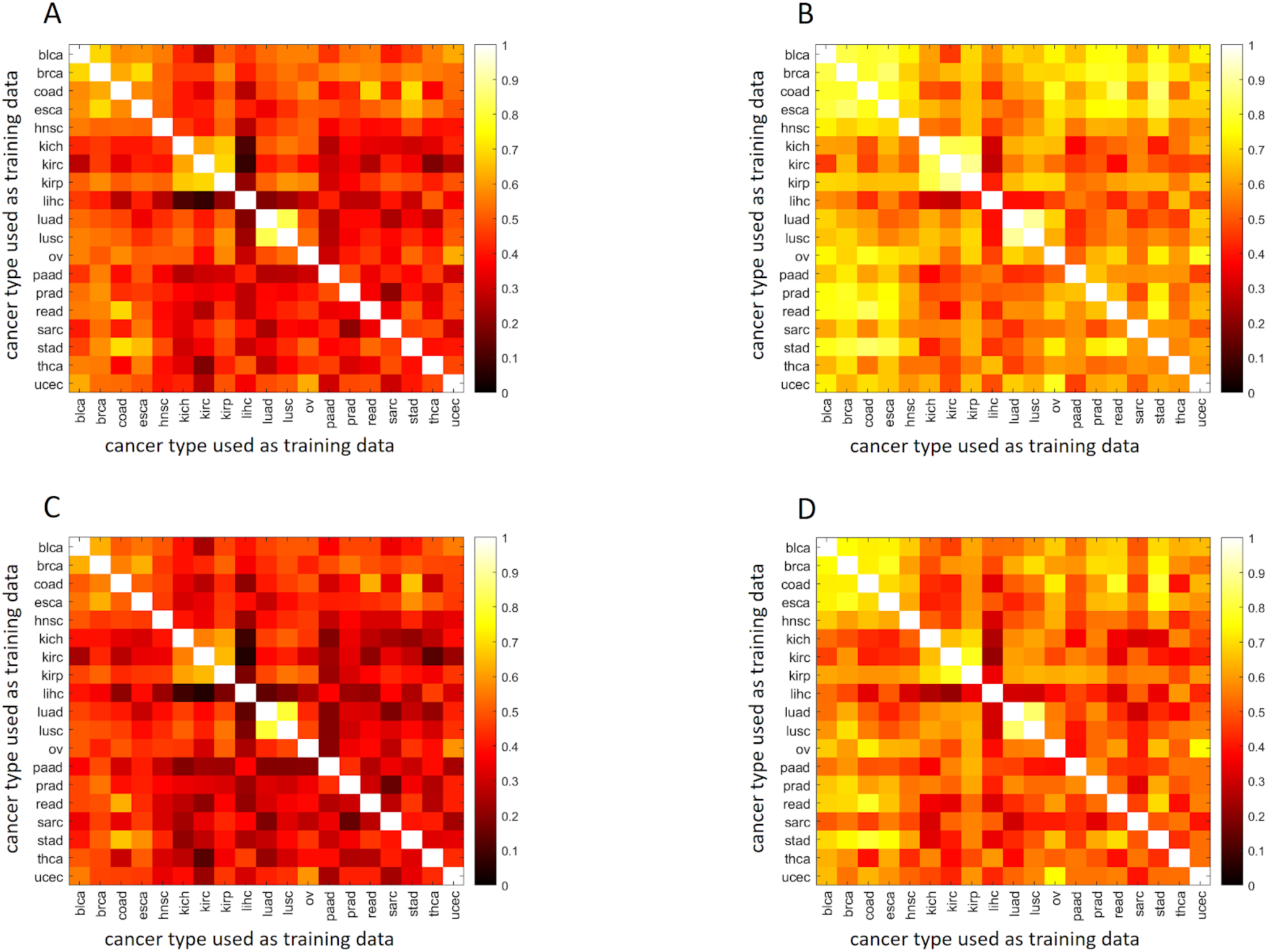
Tile level cross-classifications as a function of test set. Correlations of predicted tumor/normal status (i.e. logit of tumor probability) between pairs of classifiers, specified on the x and y-axis. Correlations are first calculated using the tile values for all slides of a given test tissue. (A) Average correlation across tissues, using both tumor and normal slides in the tissue test sets. (B) Correlation for the tissue set with the maximal correlation, using both tumor and normal slides in the tissue test sets. (C) Average correlation across tissues, using only tumor slides in the tissue test sets. (D) Correlation for the tissue set with the maximal correlation, using only tumor slides in the tissue test sets.

We hypothesized that certain tissue types might be particularly easy to classify, and to test this we tabulated which tissue sets yielded the maximal correlations for each pair of classifiers in Figure 5b (Table S2). For each pair, we listed the 3 tissue sets yielding the highest correlations. If this were random, we would expect each tissue to appear in this list 27 times. However, we observed extreme prevalence for BRCA (132 appearances, p=8.5e-119), BLCA (106 appearances, p=2.5e-43), and UCEC (62 appearances, p=1.9e-11). Many classifier pairs agree better within these 3 tissues than they do within their training tissues. Thus BRCA, BLCA, and UCEC are canonical types for intraslide spatial analysis, in addition to their high cross-classifiability at the whole slide level (Fig. 4).

### Cross-classification relationships recapitulate cancer tissue biology

To test the biological significance of cross-classification relationships, we assessed associations between tissue of origin (Hoadley et al. 2018) and cross-classification clusters. Specifically, we labeled KIRC/KIRP/KICH as pan-kidney (Ricketts et al. 2018), UCEC/BRCA/OV as pan-gynecological (pan-gyn) (Berger et al. 2018), COAD/READ/STAD as pan-gastrointestinal (pan-GI) (Liu et al. 2018), and LUAD/LUSC as lung. The hierarchical clustering in Figure 4 shows that cohorts of similar tissue of origin cluster closer together. We observed that the lung cohort clusters together on both axes, Pan-GI clusters on the test and partially the train axis, and Pan-Gyn also partially clusters on the test axis. Pan-Kidney partially clusters on both axes. To quantify this, we tested the associations between proximity of cohorts on each axis and similarity of their phenotype (i.e. tissue of origin/adeno-ness). Organ of origin was significantly associated with smaller distances in the hierarchical clustering (p-value=0.002 for test axis and p=0.009 for train axis; Gamma index permutation test. See Methods). We also grouped cohorts by adenocarcinoma/carcinoma status (Figure 4, second row from top), though SARC and ESCA do not fit either category. The cohort distances were significantly associated with adeno-ness on the test axis (p-value=0.008). We observed other intriguing relationships among cross-tissue classifications as well. Particularly, Pan-GI created a cluster with Pan-Gyn, supporting these tumor types having shared features related to malignancy. Likewise, Pan-Kidney and lung also cluster close to each other.

### Comparisons of neural networks for TP53 mutation classification

To investigate how images can be used to distinguish cancer drivers, we tested the accuracy of CNNs for classifying TP53 mutation status in 5 TCGA cohorts, namely BRCA, LUAD, STAD, COAD and BLCA. We chose these cohorts due to their high TP53 mutation frequency (Cancer Genome Atlas Research Network 2014a; Cancer Genome Atlas Network 2012; Cancer Genome Atlas Research Network 2014b), providing sufficient testing and training sets for cross-classification analysis. Using transfer learning, we obtained moderate to low AUCs for TP53 mut/wt classification (0.66 for BRCA, 0.64 for LUAD, 0.56 for STAD, 0.56 for COAD, and 0.61 for BLCA). Due to this weak performance we switched to a more computationally intensive approach in which we fully trained all parameters of the neural networks based on an architecture described in (Coudray et al. 2018) (Figure 1b), with under-sampling to address data imbalance and a 70/30 ratio of slides for training and testing. Figures 6a and 6b show heatmaps of AUC for the per-tile and per-slide classification results, respectively (see also Figures S4 and S5). Self-cohort predictions (diagonal values) have AUC values ranging from 0.65-0.80 for per-slide and 0.63-0.78 for per-tile evaluations. Stomach adenocarcinoma (slide AUC=0.65) was notably more difficult to predict than lung adenocarcinoma (slide AUC=0.80), for which we found AUC values comparable to the AUC=0.76 LUAD results reported by (Coudray et al. 2018). This LUAD fully-trained network (AUC=0.76) outperformed the transfer learning for the same data (AUC = 0.64).

**Figure 6.**
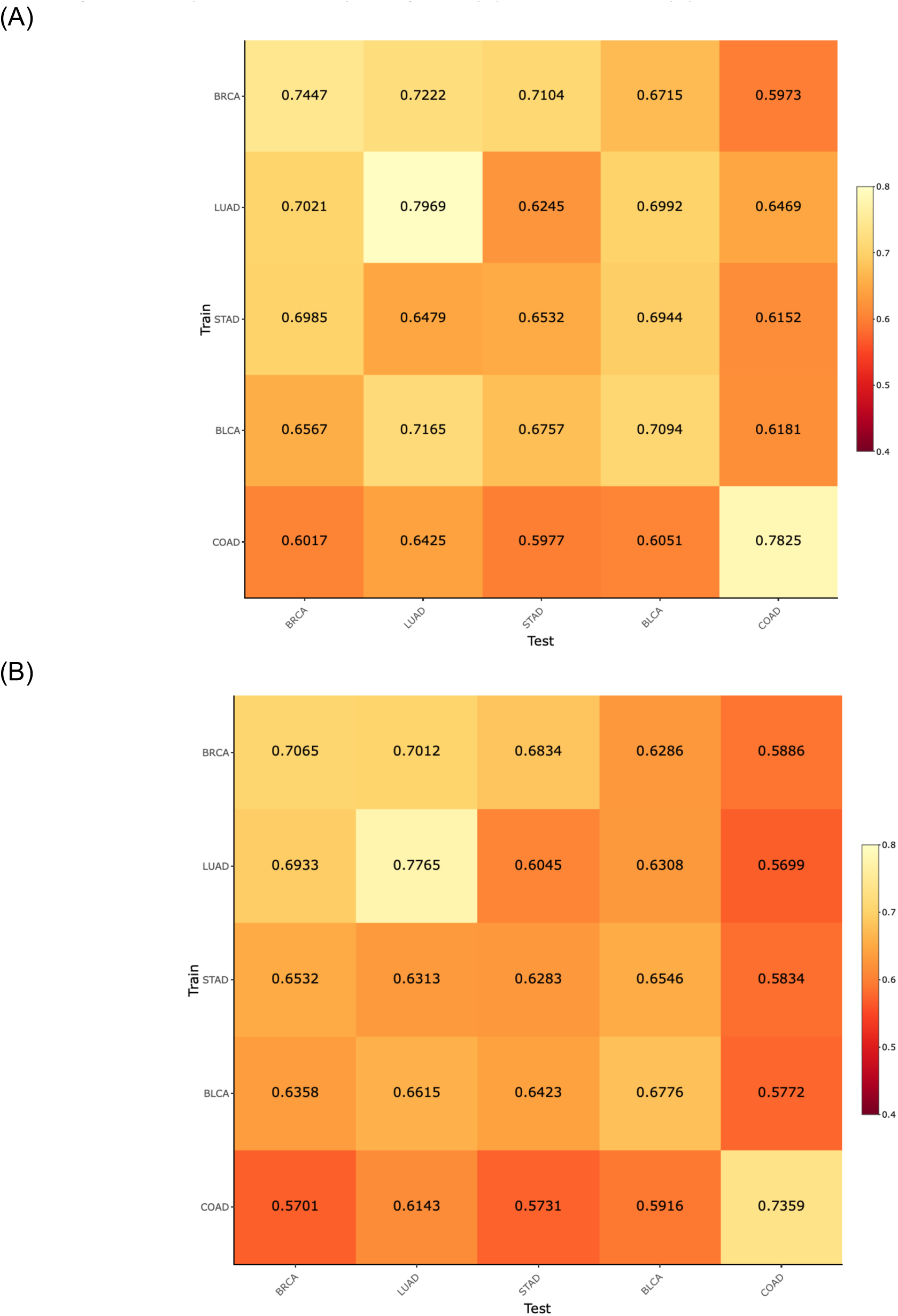
Classification of TP53 mutation status for five TCGA cohorts including BRCA, LUAD, BLCA, COAD and STAD. Cross- and self- classification AUC values from balanced deep learning models (with 95% CIs) are given (a) per-slide and (b) and per-tile.

We also tested the ability of the TP53 CNNs to cross-predict across cohorts. Cross-predictions yielded AUC values with a comparable range as the self-cohort analyses (AUCs 0.62-0.72 for slides; 0.60-0.70 for tiles), though self-cohort analyses were slightly more accurate. Colon adenocarcinoma AUC values tended to be low as both a test and train set, suggesting TP53 creates a different morphology in this tissue type. Overall, the positive cross-classifiabilities support the existence of shared TP53 morphological features across tissues. Figure 7 shows TP53 mutational heatmaps of one LUAD slide known to be mutant and one LUAD slide known to be wildtype from sequencing data. We compared the LUAD- and BRCA-trained deep learning models on these slides, as those two models provided the highest AUC values in our cross-classification experiments. Prediction maps for tumor/normal status (second row) and TP53 mutational status (third row) are shown for both samples. Both tumor/normal models correctly predicted the majority of tiles in each sample as cancer. Analogously, the BRCA-trained TP53 mutation status model predicts patterns similar to the LUAD-trained model. Importantly, the tumor/normal and TP53mut/wt classifiers highlight different regions, indicating these classifiers are utilizing distinct spatial features.

**Figure 7.**
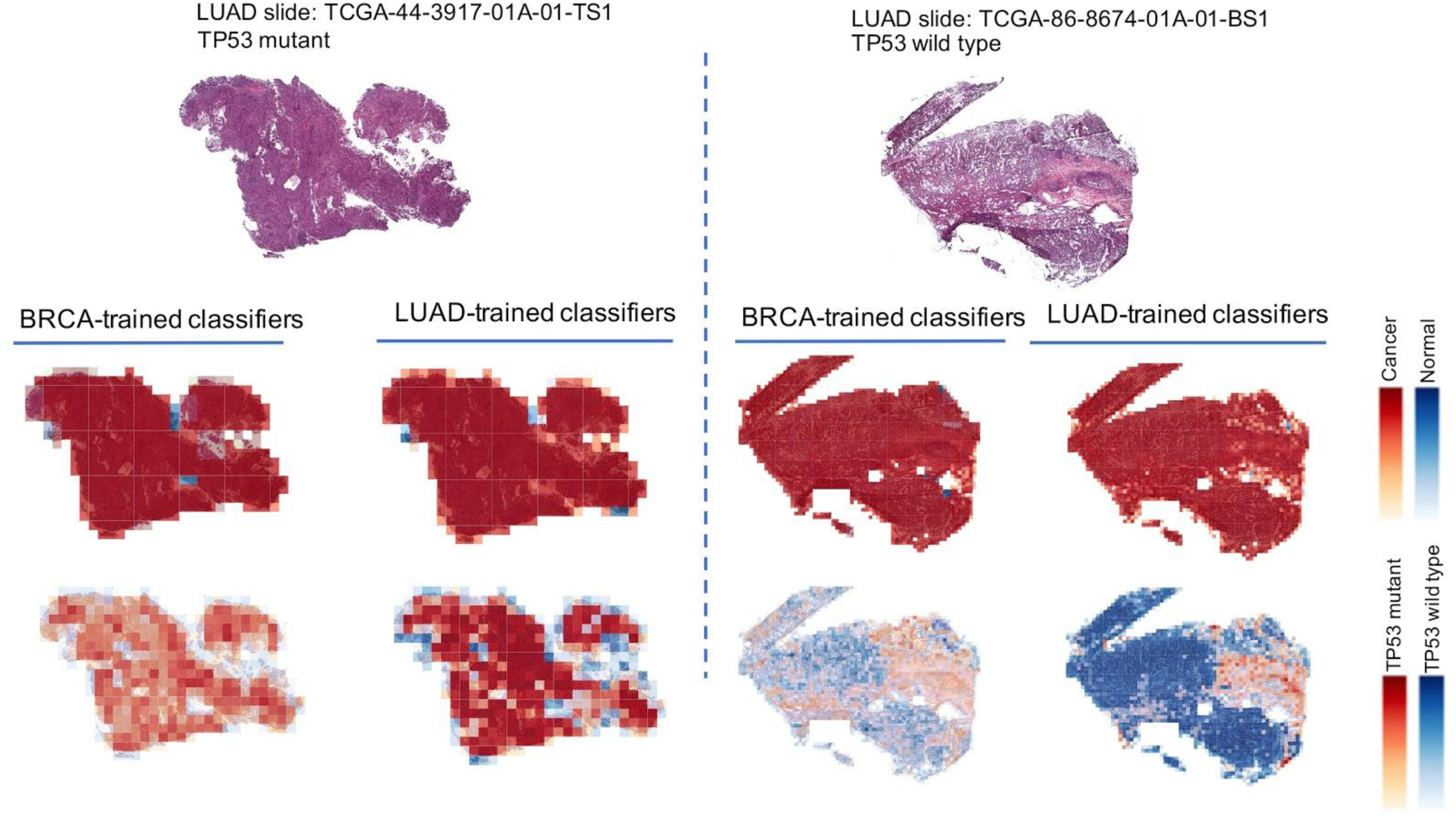
TP53 genotype heatmaps based on predicted probabilities using our deep learning model. The first row shows two LUAD H&E slides with TP53 mutant (left panel) and wild type (right panel). The second row shows prediction maps for these two slides using tumor/normal classifiers trained on BRCA and LUAD samples. Both models successfully classify samples as cancer and predict similar heatmaps. The third row shows prediction maps for these slides using TP53 mutation classifiers trained on BRCA and LUAD. The BRCA-trained and LUAD-trained heatmaps are similar, suggesting that there are spatial features for TP53 mutation that are robust across tumor types.

We next performed a tile-level cross-classification analysis as a function of test set. For most test cancer types, we observed little correlation when comparing networks trained on cancers “A” and “B” applied to test cancer “C”. Therefore, we focused on cases where C is the same as B. Figure 8 plots the correlations of TP53 mutation probability logits across cancer pairs, where each row denotes the cancer type the first CNN is trained on, and each column is both the test tissue and the second CNN training tissue. In these cases the correlation coefficients were generally positive and met statistical significance though with moderate magnitude. All correlations were significant except for the BRCA TP53 classifier applied to LUAD tumors (t-test on Fisher z-transformed correlation coefficients, FDR 5%). Notably, classifiers based on LUAD, BRCA, and COAD, worked well on BLCA, BLCA, and COAD tumors, respectively. BLCA and LUAD are the two test cancers with the largest correlations (column average). LUAD and COAD are the two training cancers with the largest correlations (row average). The high row and column averages for LUAD indicate it is canonical both as a test and a training set. Interestingly, the correlations of Figure 8 are not symmetric. For example, the network trained on LUAD achieves a correlation of 0.34 on BLCA, while the network trained on BLCA has a correlation of 0.04 when tested on LUAD.

**Figure 8.**
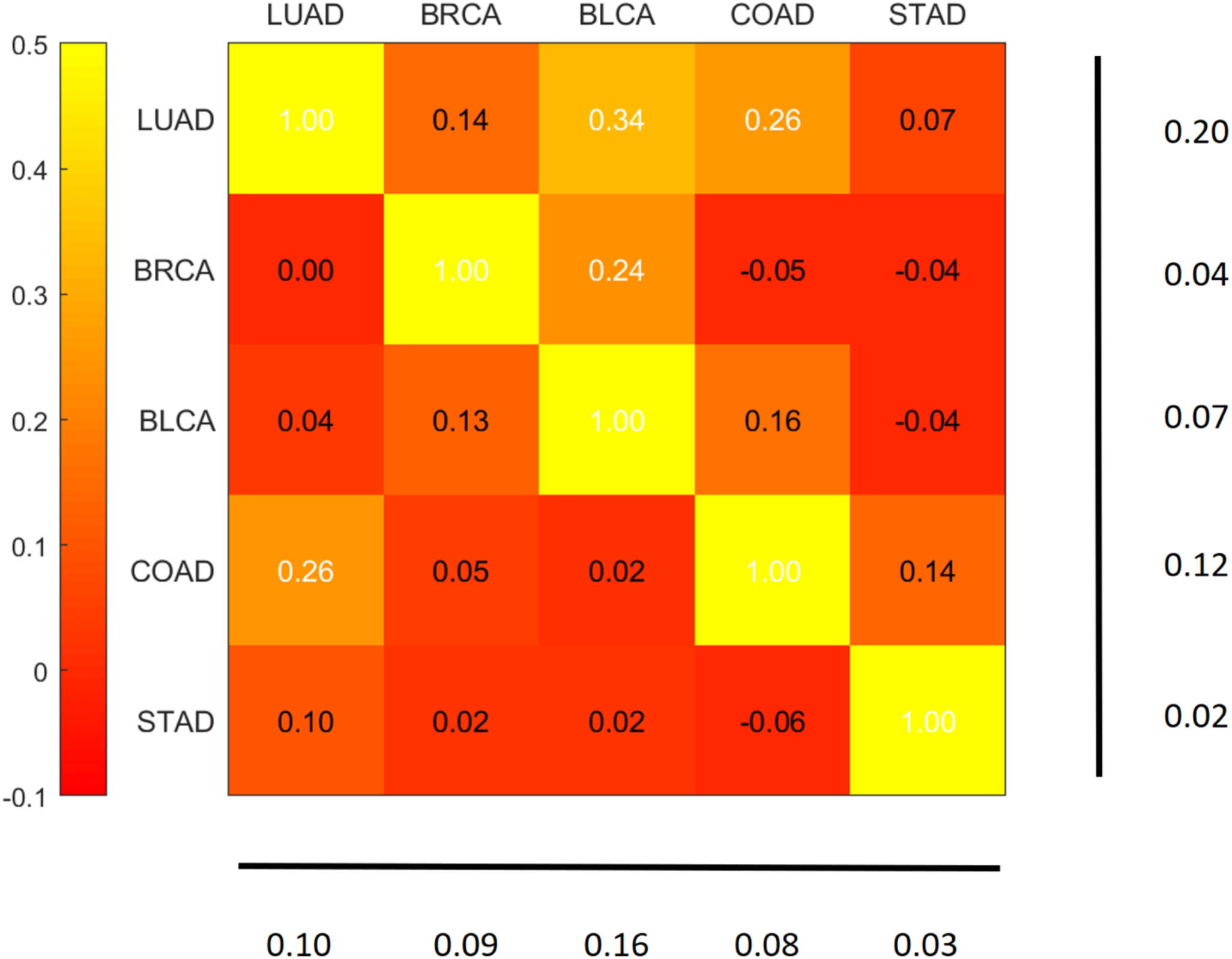
Tile-level cross classification correlations for TP53 mutational status. Row labels denote the cancer type used to train the first TP53 mutation classifier. Column labels denote both the test tissue and the tissue of the second TP53 classifier. Heatmap values indicate correlation coefficients of mutation probability logits between the two classifiers on the test tissue. Numbers at the bottom and right show column and row averages, respectively, with diagonal values excluded.

### Features impacting tumor purity prediction

TCGA provides annotations only at the whole slide level, limiting our ability to build classifiers that resolve predictive features. To better investigate features, we obtained datasets with higher resolution annotations, i.e. BreCaHAD (Aksac, et al. 2019) which provides nucleus-level tumor/normal annotations of 162 TCGA breast cancer ROIs, and 8 colorectal ROIs hand-annotated at nuclear resolution (>18,000 cells) by our group. These annotations provide exact tumor purity (the fraction of tumor cells, aka cellular malignancy) in all tiles. We trained neural networks on tile values, randomly splitting the BreCaHAD dataset into 150 *train* and 12 *test* ROIs (a total of >23,000 cells) and using the colorectal set for validation. Purities of the colorectal tiles (512×512 pixels) are spread over a wide range (mean 58%, standard deviation 19.2%), while BreCaHAD purities are higher (mean 87% in the training set), as detailed in Figure S6. These nucleus-based CNNs yielded mean absolute error of 14% and 15% for the test breast and colorectal sets, respectively. Root mean squared error (RMSE) values were 8% and 20%, respectively. Although the CNN was trained only on breast data, the average prediction for the colorectal datasets (69%) was shifted toward the true colorectal mean, suggesting that the CNN is able to learn some features common between breast and colorectal tumors.

We tested whether purity estimates were due only to local information around each cell nucleus or whether other image properties were informative. For this we first evaluated a CNN classifier (Hlavcheva et al. 2019) designed to predict tumor/normal status from individual nucleus images (see Methods). We trained on the breast nuclei, and this yielded high accuracy on reserved breast nuclei images (AUC 97-99%) However, the breast-trained CNN yielded poor classifications on the colorectal nuclei (AUC 56%), a much worse cross-classification than tile-based WSI-level analysis of TCGA data (Figure 4). We tested whether summing across nuclei within the colorectal ROIs (thousands of cells each) would improve purity predictions. However, even aggregated over full ROIs the nucleus-based RMSE was 25%, an RMSE higher than the tile-based analysis of the same data (20%). This suggests that, although the tile-based approach is not aware of individual cells, it compensates by using intercellular regions of images.

## Discussion

In this paper we have presented a versatile CNN-based framework for pan-cancer analysis of histology images. Using this framework we were able to train extremely accurate slide-based tumor/normal classifiers in nearly all cancer types, and we also were able to classify subtypes and TP53 mutation status with significant though less extreme accuracy. Critically, these pan-cancer studies enabled us to compare classifier outputs as a function of training tissue, test tissue, and neural network architecture. We found that many tumor images have robust intra-slide structure that can be consistently identified across CNN classifiers. Our findings can be viewed in three prongs: identification of pan-cancer morphological similarities, transfer learning as a common feature extractor, and interpreting spatial structures with tumors.

### Identifying pan-cancer morphological similarities

While other recent works have investigated image-based cancer classification (Fu et al. 2019; Kather et al. 2019), cross-classification has until now been little studied. Comparisons of classifiers support the existence of morphological features shared across cancer types, as many cross-cancer predictors achieve high AUCs. Specific relationships between types are also informative. Cancers from a common tissue, such as (KIRC, KIRP, KICH), (LUAD, LUAC), and pan-GI cancers are good predictors of each other, and there are also significant similarities within adenocarcinomas and carcinomas, respectively. Remarkably, BRCA, BLCA, and UCEC are unexpectedly easy to classify as test sets, showing strong cross-classifiability both at the WSI and tile levels. Further studies are likely to benefit from focusing on these as canonical image types for analysis and method development. Interestingly, this behavior is not symmetric between train and test cancer types. For example, while the network trained on KIRC achieves an AUC >90% when tested on BLCA, training on BLCA and testing on KIRC results in an AUC <65%.

A next challenge is to better define the morphological features that underlie cross-classifiability. One approach would be to select classifiers with highly similar outputs and then overlap their spatial salience maps (Yosinski et al. 2015; Samek et al. 2017). Another approach would be to assess if shared morphological features are predictive of shared genomic markers, e.g. via nonlinear canonical correlation analysis (see Lai and Fyfe 2000 and Hsieh 2000 for examples). For these types of image morphology questions, careful selection of cancer test sets will be critical. For example, for the TP53 mutation studies we had enough data to identify significant cross-correlations and spatial structures within images, but such analysis will be more challenging for rarer drivers.

### Transfer learning as a common feature extractor

Transfer learning-based methods use a universal set of pre-trained layers based on non-histological image collections to decompose images into features, an aspect which reduces computational costs but can limit classification accuracy. While transfer learning networks excel at tumor/normal classification, they have lower accuracy for cancer subtype and mutation status predictions. This may be because the features associated with subtype and mutation status are not well-represented in the non-histological image collections. We found that fully trained models, which learn all network parameters directly from the cancer images and are computationally more demanding, yielded higher AUCs. Thus the suitability of transfer learning is task-specific, though determining which tasks are suitable is an open challenge. The effectiveness of CNN architectures can also be impacted by class imbalances in the histopathology samples, an issue that would be further exacerbated by intratumoral heterogeneity. This general problem corresponds to multi-label, multi-instance supervised learning with imbalanced data, an active area of machine learning research (Charte et al. 2015; Read et al. 2015; Li and Wang 2016).

### Interpreting spatial structures within tumors

Cross-comparisons of classifiers can highlight robust spatial structures within tumor images (e.g. Figure 8), but interpretation remains a major challenge. Neural networks provide only indirect information about the features responsible for such structures, and expert manual pathological analysis of such cases will be essential. Manual analyses may also clarify the identity of predictive features whose existence is supported by CNNs. For example, our comparison of tile and nucleus-level approaches indicated that intercellular regions are useful in predicting tumor purity, but it is uncertain what specific features mediate this relationship. It is worth noting that such analyses would not be possible without mixtures of tumor and normal regions together within images. Thus it will be important to analyze regions with spatial diversity rather than only regions of high purity, which has been the focus of some recent works (Fu et al. 2019). Finally, the field would be advanced if fine-grained spatial pathological annotations can be generated at scale by the community, e.g. through further curation of TCGA and other cohorts. Prior single histology studies have distinguished spatially important regions by training on detailed annotations from pathologists (Wei et al. 2019), and expansion across histologies would enable further understanding through cross-comparisons of classifiers. Such comparisons of genetically and phenotypically diverse tissues will be a potent approach to reveal morphological structures underlying cancer biology.

## Materials and Methods

### Transfer learning

#### Sample selection for tumor/normal classification

Since there are very few normal FFPE WSIs on TCGA, we only considered flash frozen samples (with barcodes ending with BS, MS, or TS). We selected 19 TCGA cohorts that had at least 25 normal samples. The samples were randomly divided into 70% training and 30% testing. Stratified sampling was used to balance the ratio of positives and negatives into train and test sets.

#### Sample selection for subtype classification

WSI images from 10 tissue types were used for subtype classification. FFPE and flash frozen samples are approximately balanced among the tumor WSIs; hence we used both for subtype classification. The samples were randomly divided into 70% training and 30% testing. Some cancer tissues had subtypes that were available as individual cohorts within TCGA. These 3 tissues were LUAD/LUSC (lung); KICH/KIRC/KIRP (kidney); and UCS/UCEC (uterine). For all other tissues, TCGA provided single cohorts that spanned multiple subtypes designated by pathologist annotations. Only clinical subtype annotations with at least 15 samples were considered. Samples with ambiguous or uninformative annotations were not included.

#### CNN architecture and training

We used a Google Inception v3-based architecture for pan-cancer tumor/normal classification of TCGA H&E slides. Our CNN architecture uses transfer learning on the Inception module with a modified last layer to perform the classification task. For predicting mutational status we utilize the same architecture as in Coudray et.al. (Coudray et al. 2018) and fully trained the model on TCGA WSIs.

The output of the last fully connected layer of Inception v3 (with 2048 neurons) was fed into a fully connected layer with 1024 neurons. The output was encoded as a one-hot-encoded vector. A softmax function was utilized to generate class probabilities. Each training simulation was run for 2000 steps in batches of 512 samples, with 20% dropout. Mini-batch gradient descent was performed using Adam optimizer (Kingma and Ba 2014). To mitigate the effects of label imbalance in tumor/normal classification, undersampling was performed during training by rejecting inputs from the larger class according to class imbalances, such that, on average, the CNN receives equal number of tumor and normal tiles as input. Per-tile classification ROCs were calculated based on thresholding softmax probabilities and per-slide classification ROCs were based on voting on maximum softmax probability. Due to significant additional compute costs we did not optimize on hyper parameters, e.g. number of epochs or learning rate, instead using common values for similar image classification problems. Details are in the Github code repository.

#### Preprocessing and transfer learning steps

1. Aperio SVS files from primary solid tumors or solid tissue normal samples with 20X or 40X magnification were selected.
2. Each SVS file was randomly assigned to train or test set.
3. 40X images were resized to 20X.
4. Background was removed as in (Coudray et al. 2018).
5. Images were tiled into non-overlapping patches of 512×512 pixels.
6. Tiles were used as inputs of the Inception v3 network (pretrained on ImageNet; downloaded from http://download.tensorflow.org/models/image/imagenet/inception-2015-12-05.tgz), in a forward pass and the values of last fully connected layer (‘pool_3/_reshape:0’) were stored as ‘caches’ (vectors of 2048 floating point values).
7. Caches from similar holdout group were shuffled and assigned to a TFRecords in groups of 10,000.
8. TFRecords were used as input to the transfer learning layers.

#### Programming details

All analysis was performed in Python. Neural network codes were written in TensorFlow (Martín et al. 2016). Images were analyzed using OpenSlide (Goode et al. 2013). Classification metrics were calculated using Scikit-learn (Pedregosa et al. 2011). All transfer learning analysis including preprocessing was performed on the Google Cloud Platform (GCP). The following GCP services were used in our analysis: Kubernetes, Datastore, Cloud Storage, and Pub/Sub. During the preprocessing steps we used up to 1,000 compute instances (each 8 vCPUs and 52GB memory) and up to 4,000 Kubernetes pods. Cloud Storage was used as shared storage system, while Pub/Sub asynchronous messaging service in conjunction with Datastore were used for task distribution and job monitoring of the Kubernetes cluster. This architecture ensures scalability and a fault tolerant process. We leveraged a similar architecture for the pan-cancer training/testing process.

### Mutational classification

#### Sample selection for mutational classification

We selected flash frozen WSIs of BRCA, LUAD and STAD cohorts. Impactful TP53 mutations were determined using masked somatic mutations maf files called by MuTect2 (Cibulskis et al. 2013). Mutations which were categorized as MODERATE/HIGH (by VEP software (McLaren et al. 2016)) in the IMPACT column were considered as impactful mutations. If the gene had at least one impactful mutation in the sample, it was counted as mutated and was considered as wild-type otherwise. Table 1 shows the number of wild type and mutated slides in each cancer type. For cross classification, the model was trained on the entire training cohort and predictions were made on the entire test cohort.

**Table 1.**
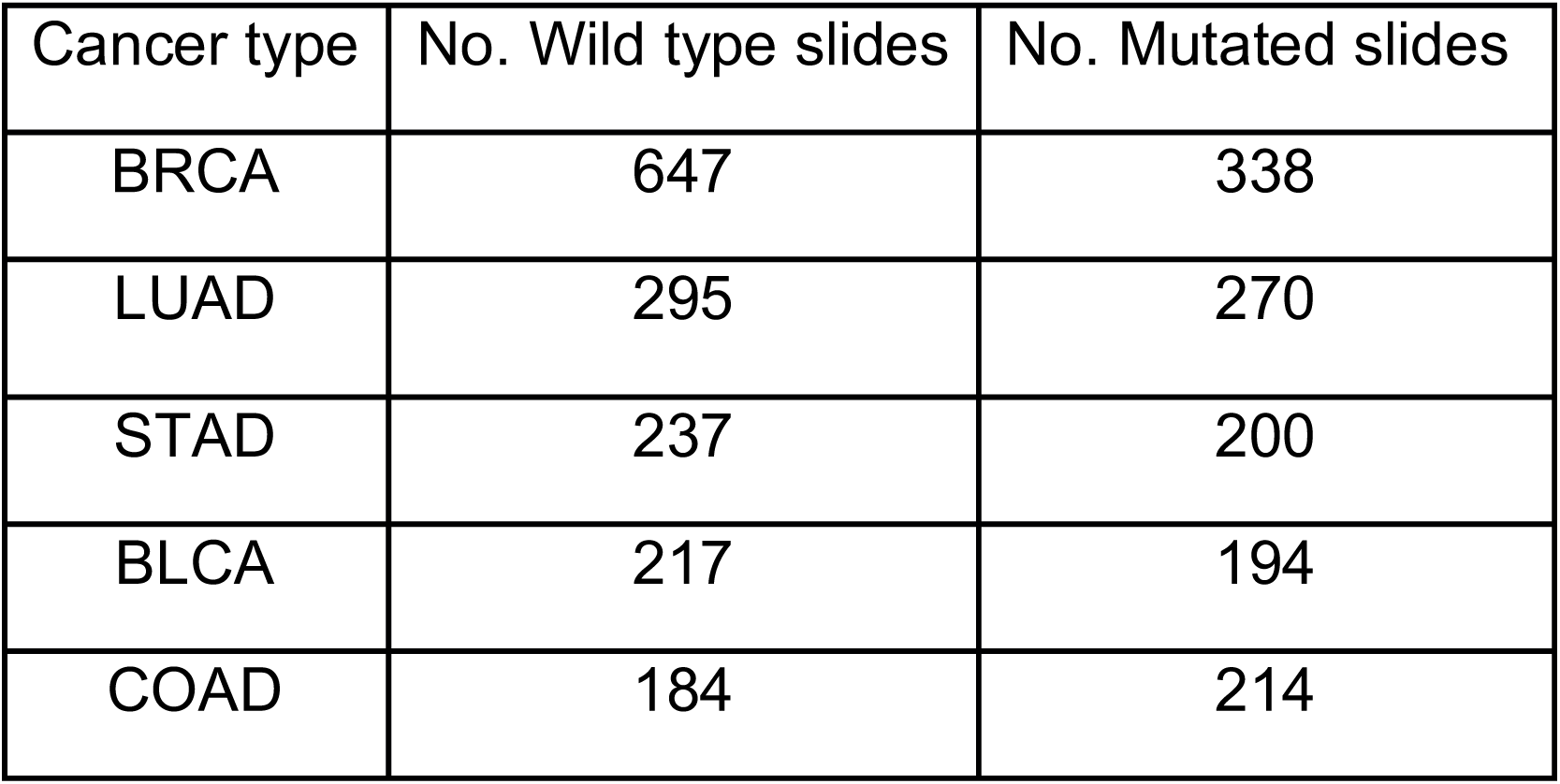
Numbers of wild type and mutated slides in each TP53 cohort.

| Cancer type | No. Wild type slides | No. Mutated slides |
| --- | --- | --- |
| BRCA | 647 | 338 |
| LUAD | 295 | 270 |
| STAD | 237 | 200 |
| BLCA | 217 | 194 |
| COAD | 184 | 214 |

#### CNN architecture and training

We utilized the Inception v3 architecture (Coudray et al. 2018) to predict TP53-associated mutations in BRCA, LUAD and STAD cohorts. Unlike the tumor/normal analysis, transfer learning was not used for mutational classifiers. Instead models were fully trained on input slides. As a pre-processing step, we used a fully trained normal/tumor classifier to identify and exclude normal tiles within each tumor slide. This filtering step ensures that tiles with positive mutation class label are also labeled as tumor. To predict mutations in the TP53 gene, we trained 2-way classifiers, assigning 70% of the images in each tissue to training and the remaining 30% to the test set. Cross-tissue mutational classification was performed by training the model on the entire train set of a cohort and performing prediction on other cohorts. The model outputs for tiles were used to produce slide level prediction by averaging probabilities. A similar downsampling as in the tumor/normal classifier was performed to handle data imbalance issues.

#### Computational configuration

All of the computational tasks for mutation prediction were performed on linux High performance computing clusters with following specification: 8 CPUs, RAM: 64 GB, and Tesla V100 GPUs, 256 GB RAM. Furthermore, The GPU-supported TensorFlow needed CUDA 8.0 Toolkit and cuDNN v5.1. All GPU settings and details were obtained from TensorFlow and TF-slim documentations and NVIDIA GPUs support.

#### Cross-classification statistics

Hierarchical clustering was applied to cross-classification per-slide AUC values using UPGMA with Euclidean distance. To determine the association between clustering and independent phenotypic labels (i.e. organ and adeno-ness), we used Gamma index of spatial autocorrelation from the Python package PySal (Rey and Anselin 2007). Gamma index is defined as (Hubert, Golledge, and Costanzo 1981):

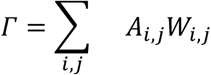

where *A* is the feature matrix and *W* is the weight matrix, and indices range over cancer cohorts. For each axis and each phenotype group (i.e. organ or adeno-ness), we calculate a separate Gamma index. We define *A*_*i,j*_ = 1 if cohorts *i* and j have the same phenotype (e.g both are adenocarcinoma) and *A*_*i,j*_ = 0 otherwise. For weights we set *W*_*i,j*_ = 1 if cohorts *i* and *j* are immediately clustered next to each other and *W*_*i,j*_ = 0 otherwise. P-values are then calculated by permutation test using the PySal package. We dropped any cohort with ‘Other’ phenotype from this analysis. To avoid extensive computation cost for computing CIs, we used the method of Reiser (2000) to compute CIs instead of generating bootstrap subsamples. A similar procedure is used to compute the CIs of tumor/normal and cancer subtype classifiers. In order to compute tile level correlations we first compute tumor probability logit for each tile, defined as log((p+*ε*)/(1-p+*ε*)), where p is tumor probability and *ε*=0.0001 is added to avoid dividing by or taking logarithm of zero.

### Purity estimation for BreCaHAD and Colorectal nuclear annotations

#### Tile-based purity estimation

We used Inception, DenseNet, and Xception-based transfer learning models, each trained for 20 epochs, where the network at the epoch >10 performing best on test data and having test mean squared error larger than the train set is used for validation. Tiles of size 128×128 resulted in large test errors, and the ROIs were too small for 1024×1024 tiles. We therefore focused on tiles of sizes 512 and 256, and tiles of size 512 for validation. For cases with reduced magnification we down-sampled 512-by-512 tiles by a factor of two. To correct for acquisition differences between breast and colon cancer ROIs, we equalized the tile histogram distribution. For each tile, purity is defined as the ratio of tumor cells to total cells. We slightly adjusted to avoid purities too close to zero or one, as these may destabilize the analysis, i.e. given a tile with purity value p, we compute logit purity as log((p+0.05)/(1.05-p)), then invert the logit to obtain adjusted purity values. We used overlapping tiles with step size 64 pixels for both tile sizes. Given the extracted features, we used a fully connected layer of 256 neurons with ReLU non-linearity, followed by a drop out of 25%, and a fully connected neuron using the sigmoid activation. We used the “he_normal” initialization method of Keras described in He et al (2015), and an elastic net regularization setting L1 and L2 penalties to 0.0001.

#### Nucleus-based purity estimation

We implemented the network of Hlavcheva et al. (2019) including their reported hyperparameters. The goal of this method is to classify individual nuclei as tumor or normal. Nucleus patches were resized to 32 by 32 pixels. To adjust for acquisition differences between the breast and colon datasets we applied histogram equalization to both datasets. We trained on the BreCaHAD training set and tested on the reserved breast data across all individual nuclei, finding high accuracy (AUC 97-99%). For comparison, we also tested a transfer-learning approach. The transfer learning pipeline used similar preprocessing, except nucleus patches were resized to 128 by 128 pixels since Inception requires images to be larger than 75 by 75 pixels. The fully-trained method was superior to transfer-learning (all transfer-learning AUCs < 65%, over various parameter choices). Therefore for analysis of the colon cancer dataset, we used the Hlavcheva et al fully-trained method, trained on the entire BreCaHAD dataset. For predictions of TPF on ROIs, we compared the sum of predicted tumor probabilities across all nuclei to the pathologist purity annotations of all cells in the ROI.

## Data/Code Availability

Code used in this analysis can be found on the GitHub page: https://github.com/javadnoorb/HistCNN.

## Acknowledgements

This research benefited from the use of credits from the National Institutes of Health (NIH) Cloud Credits Model Pilot, a component of the NIH Big Data to Knowledge (BD2K) program. This material is based upon work supported by Google Cloud. JHC acknowledges support from NCI grant R01CA230031. JN and JHC would like to thank Yun-Suhk Suh for helpful comments.

## Author Contributions

JN and SF developed the TCGA CNN implementations, analyzed the TCGA data, and drafted the manuscript. AF developed and analyzed the CNN implementations for the nuclear data, contributed to the analysis of TCGA data, and drafted the manuscript. DC and DR developed the nuclear cell datasets, contributed to data analysis, and provided pathological evaluations. MS developed the cloud software engineering approaches and contributed to data analysis and drafting of the manuscript. KZ oversaw the methods development and statistical analysis and drafted the manuscript. JHC led the project and finalized the manuscript.

## Supplementary Figures

**Figure S1.**
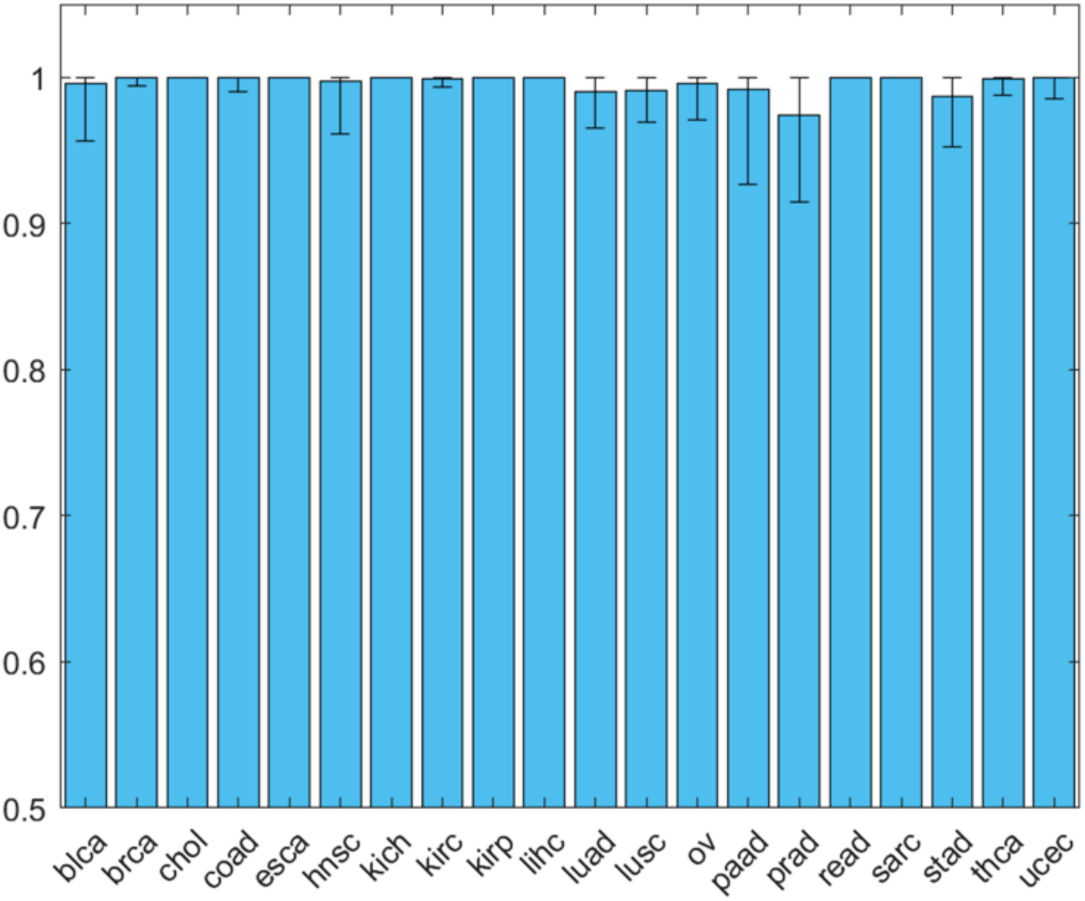
Per-slide AUC values for tumor/normal classifiers and with their confidence intervals where similar tumor types are used for training and testing. The height of each bar denotes the mean AUC and error bars denote the lower and upper bounds of the CI. Cancer types with small or imbalanced test data tend to have larger and skewed CIs. Note that a generic CI cannot be assigned to cancer types with an AUC of 100%.

**Figure S2.**
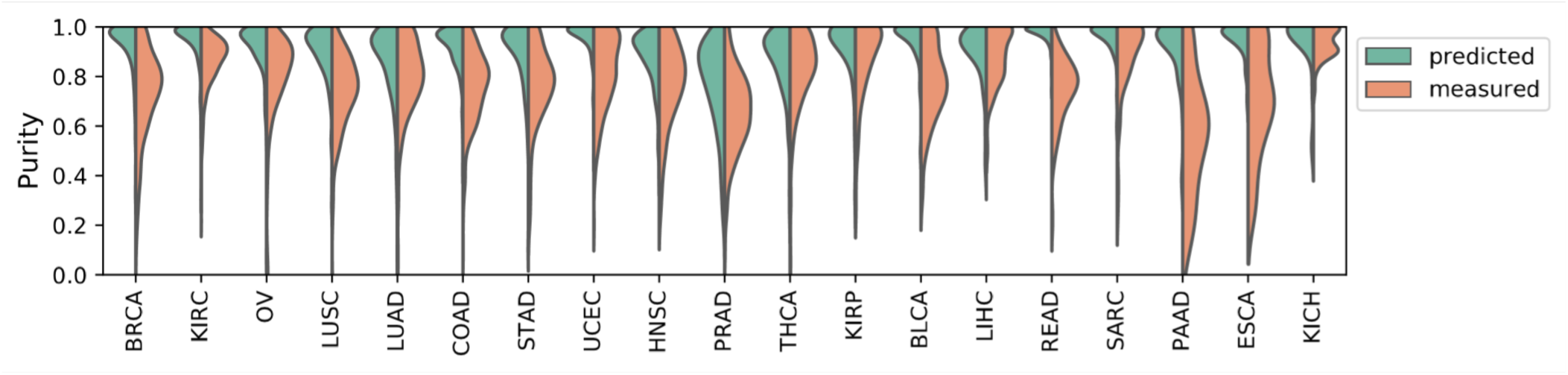
Distribution of tumor purity as predicted by our CNN model compared to the pathologist reports.

**Figure S3.**
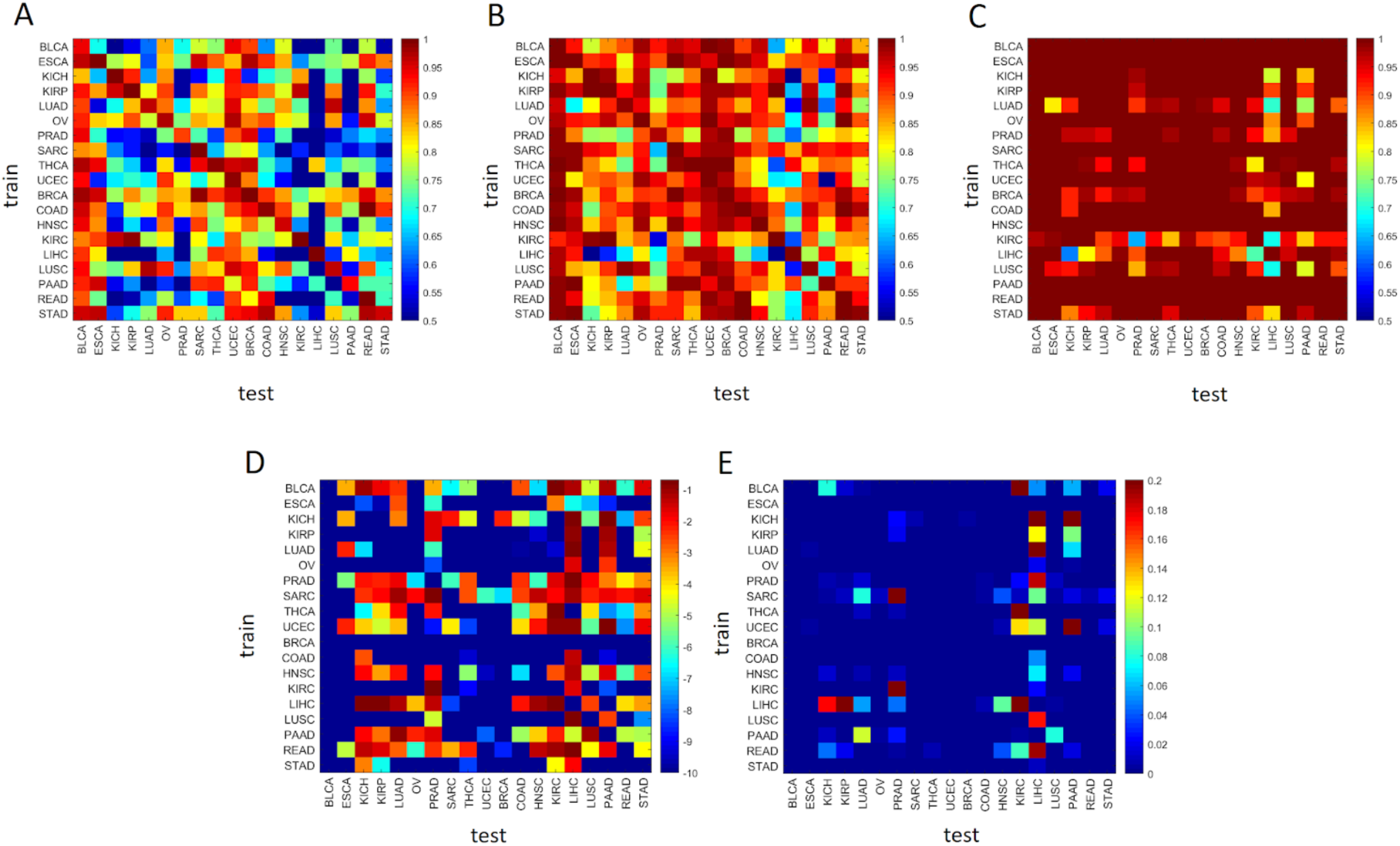
Confidence interval of AUCs for tumor normal and cross classification models. The lower bound of the CI, mean AUC, and upper bound of the CI are presented in subfigures (A), (B), and (C), respectively. Out of the 19*19=361 cross-classification models, the lower bound on the CI of 164 models is above 80%, suggesting the presence of strong common morphological features across various cancer types. Subfigures (D) and (E) provide the log10 and adjusted p-values of the hypothesis tests for AUCs being larger than 0.5 (null AUC=0.5, alternative AUC>0.5). 330 out the 361 classification models are significant (have AUC>0.5) while bounding FDR by 5%.

**Figure S4.**
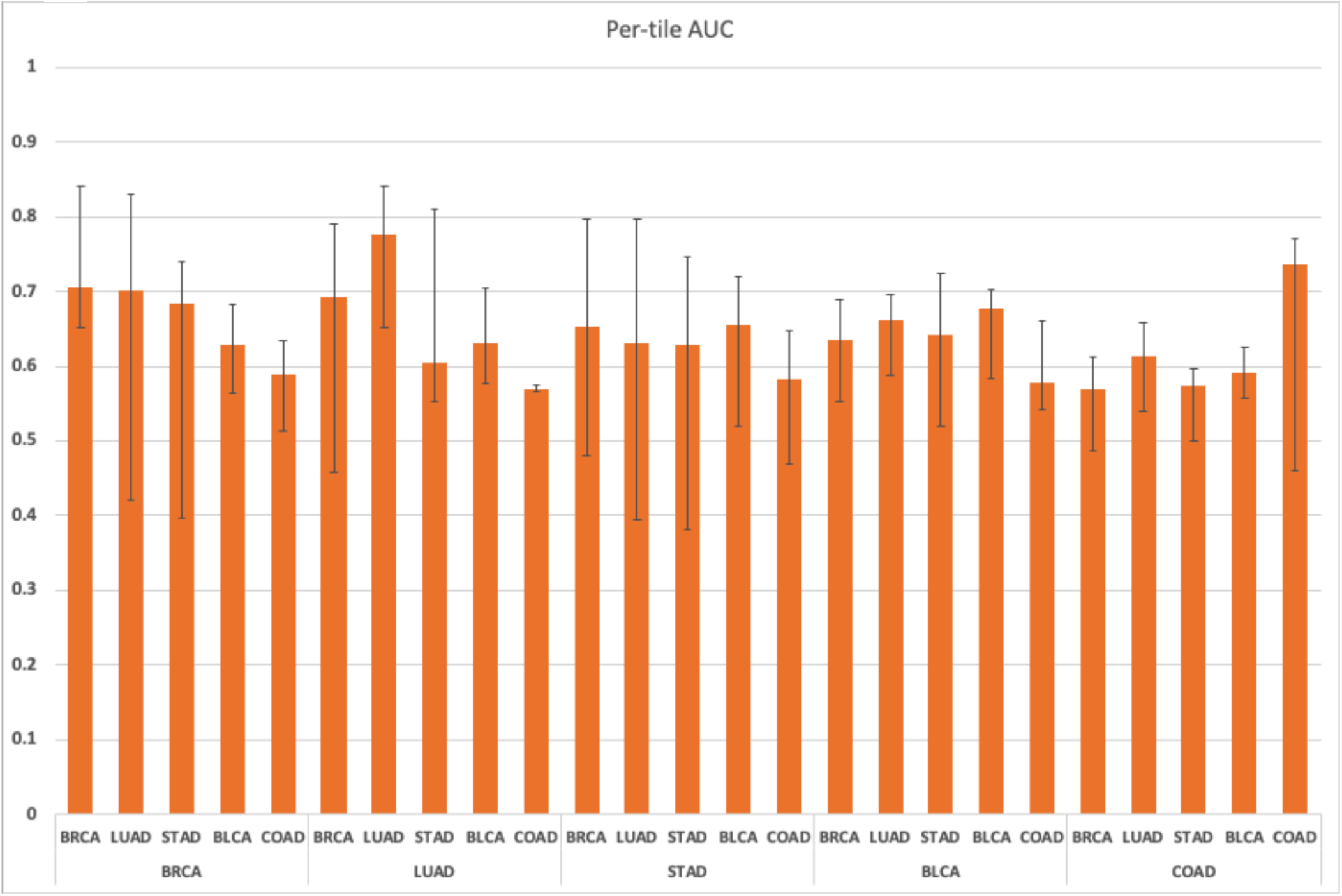
Per-tile AUC values for TP53 mutational status cross-classification experiments along with their confidence intervals

**Figure S5.**
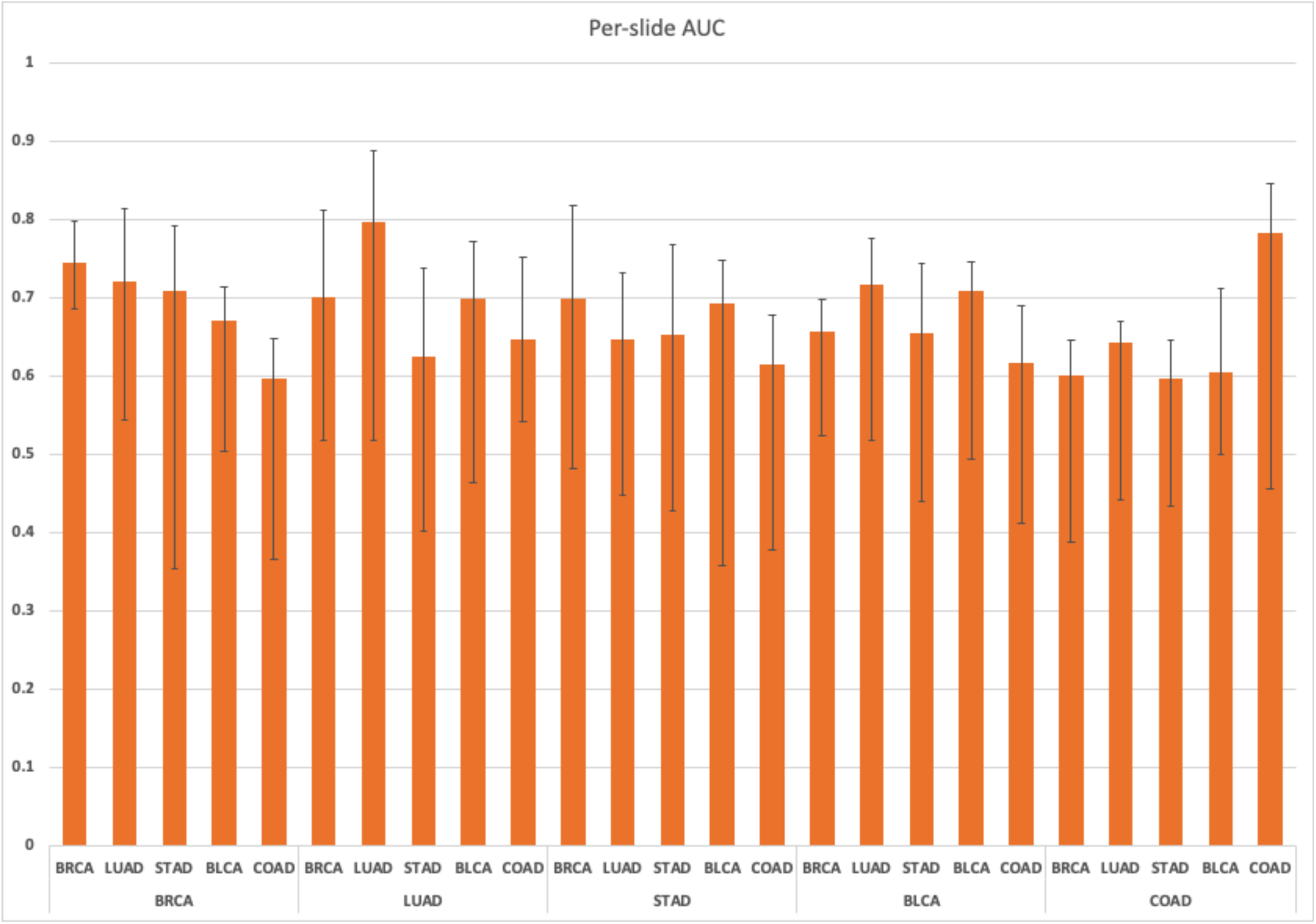
Per-slide AUC values for TP53 mutational status cross-classification experiments along with their confidence intervals

**Figure S6.**
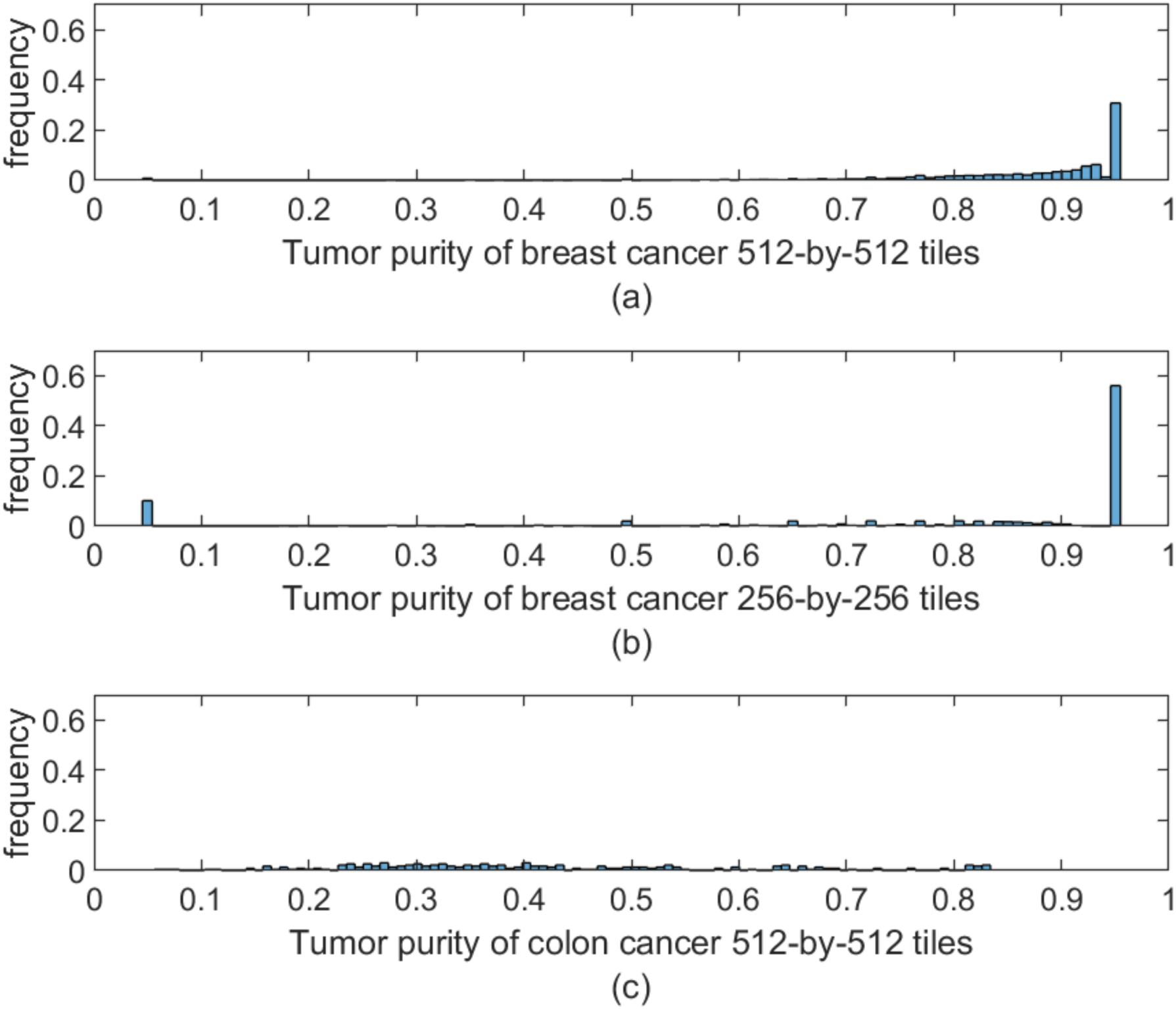
Purity histogram of breast and colon cancer ROIs. Distribution of regularized TPF across the tiles of breast and colon cancer ROIs. While the breast cancer dataset used for training is mostly comprised of tiles with large TPFs, the colon cancer validation cohort has a more spread TPF.

**Table S1**. AUC and its corresponding CI for cancer subtype classification at the slide level. P-values are based on one versus all comparisons, null expectation AUC=0.5.

**Table S2**. Cross-classifier correlations for the 3 test tissues with the maximal correlations.

## Notes

#### Summary of Updates

The new version is a significant rewrite emphasizing novel cross-classification analyses and adding a new section on comparative analysis of images at nuclear resolution. Furthermore the TP53 mutational classifications are expanded to more cancer types and individual tumor maps are included and compared.

